## Supplementary material for "Fluctuations in first passage times and utility of resetting protocol in biochemical systems with two-state toggling": Manuscript File

### Non-monotonicity of fluctuation of FPT, starting from the “+” state

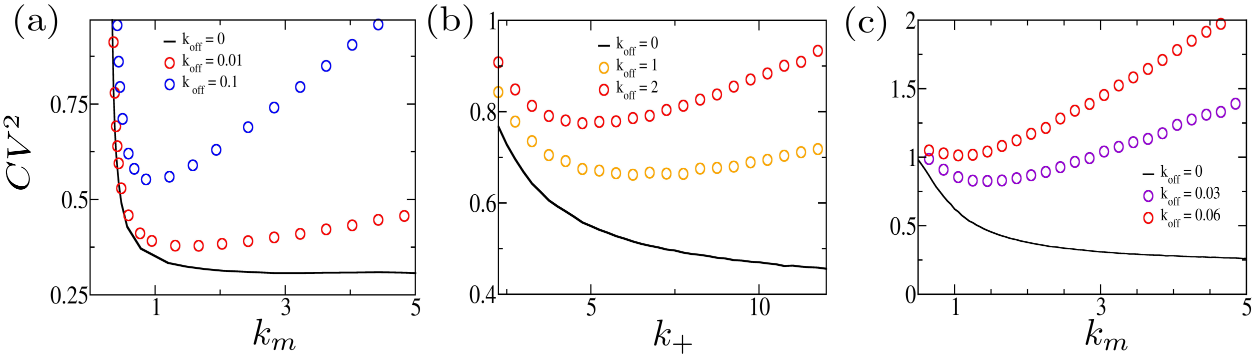


Figure S1: $CV^2$ of FPT for the initial state being “+”, is plotted against the bias strength of the three models studied in the main manuscript. In all the graphs, the case of $k_{off} = 0$ (no toggling), is shown in solid black line – we see that there is no non-monotonic behaviour of $CV^2$. With state toggling and the rise of $k_{off}$, the fluctuations rise and the U-shape is seen. (a) In the population dynamics model, the bias is the death rate per cell $k_d$. Here $k_b = 0.3$, $n_0 = 5$, and $k_{on} = 0.51$.(b) For the membrane-ECM adhesion model, the bias strength is the detachment rate constant $k_+$. Here $k_{on} = 20$, $F = 3.5$, $N_0 = 3$, $\bar{N} = 10$, $k_0 = 0.2$, $\gamma = 1$,and $f_d = 1$. (c) In the regulated gene transcription model, the bias is the transcription rate $k_m$. The parameters used are $\gamma = 0.1$, $k_{on} = 0.2$, and $X = 10$.
